# Reproductive resilience but not root architecture underpin yield improvement in maize (*Zea mays* L.)

**DOI:** 10.1101/2020.09.30.320937

**Authors:** Carlos Messina, Mark Cooper, Dan McDonald, Hanna Poffenbarger, Randy Clark, Andrea Salinas, Yinan Fang, Carla Gho, Tom Tang, Geoff Graham

## Abstract

Plants capture soil resources to produce the grains required to feed a growing population. Because plants capture water and nutrients through roots, it was proposed that changes in root systems architecture (RSA) underpin the three-fold increase in maize grain yield over the last century^1,2,3,4^. Within this framework, improvements in reproductive resilience due to selection are caused by increased water capture^1^. Here we show that both root architecture and yield have changed with decades of maize breeding, but not the water capture. Consistent with Darwinian agriculture^5^ theory, improved reproductive resilience^6,7^ enabled farmers increase the number of plants per unit land^8,9,10^, capture soil resources, and produced more dry matter and grain. Throughout the last century, selection operated to adapt roots to crowding, enabling reallocation of C from large root systems to the growing ear and the small roots of plants cultivated in high plant populations in modern agriculture.

---

In the U.S., crop improvement in temperate maize has resulted from pedigree breeding and reciprocal recurrent selection^9,12^, and the optimization of agronomic practices such as planting density ^9,10,12^. In the 1920s, breeders used double crosses (DX) to economically produce seed. This process was replaced by single hybrids (SX) in the 1950s, when more productive inbred lines resulted from the breeding efforts. The finding that genetic gain is more pronounced at higher plant populations ^9,11^ suggests that modern maize genotypes have increased tolerance to stress. The greater stress tolerance may be attributed to increased resource capture and/or enhanced reproductive resilience, but the relative importance of these two factors is unknown.

A common plant adaptation to cultivated systems is the reduction of intraspecific competitive ability^5^. Because of genetic segregation and lower cycles of selection, higher intraspecific competition, emergence of stratified plant sizes, and low yielding dominated plants within a population^13^ is expected in DX, but to a much lower degree in SX. The more genetically and phenotypically uniform SX germplasm can produce deeper and more uniform root systems than DX, where small plants contribute root mass only in the top soil horizons. This population-emergent phenotype on RSA can influence patterns of water uptake in a manner consistent with the hypothesis proposed by Hammer et al. (2009)^1^, which states that changes in root angles in response to selection for yield, underpin increased water capture and the yield response to plant population. This hypothesis is consistent with the observation that canopy temperature decreased with increasing year of commercialization in a set of hybrids grown under water deficit^14^, and simulation of breeding strategies for improved drought tolerance^2^. Simulations of plant populations with contrasting plant-to-plant variation in RSA shows increased root length density in the top soil horizons in DX relative to SX (Fig. 1a,b).

**Figure 1.**
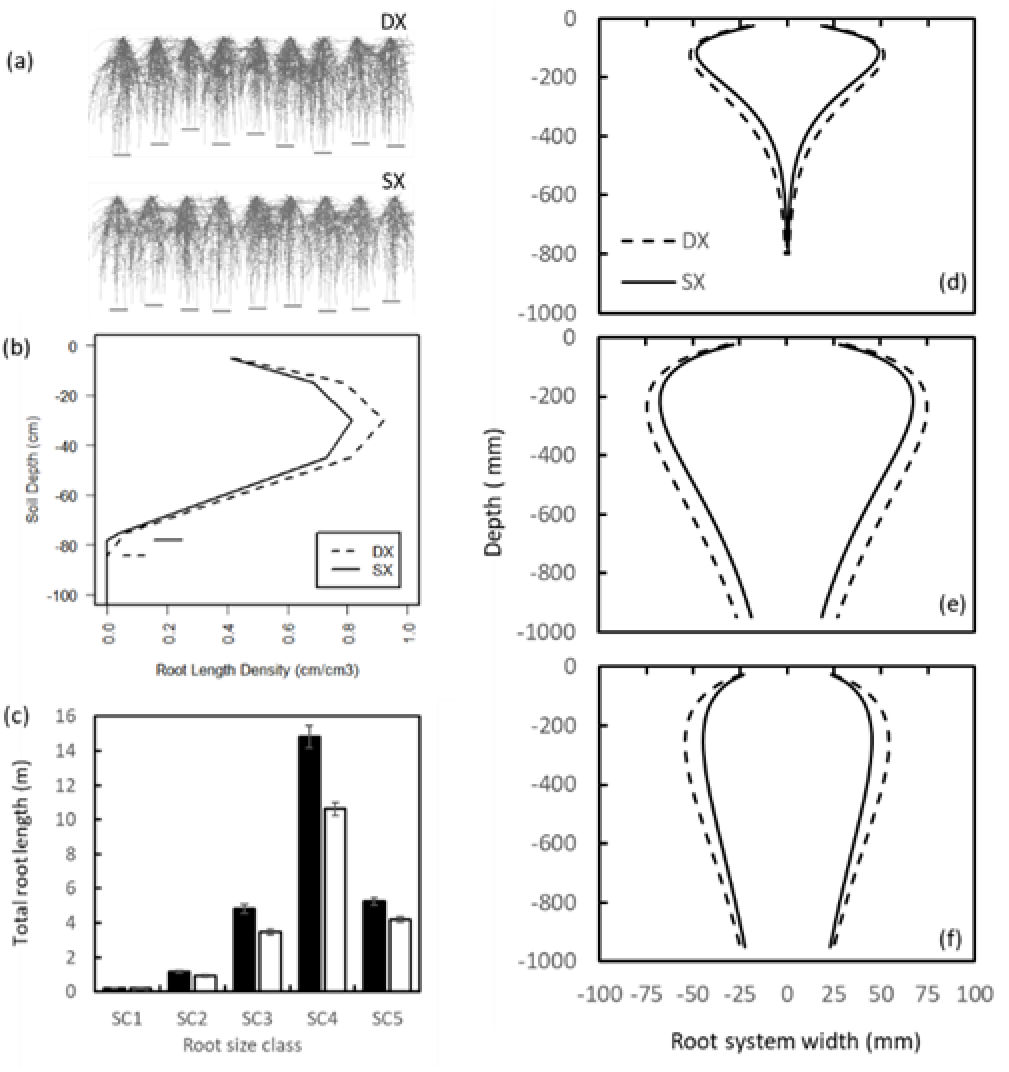
Simulated and observed root systems. A) Simulated architecture for double (DX) and single cross (SX) maize hybrids accounting for plant-to-plant variation in size using a Corteva proprietary software and visualized using ParaView (Kitware, Clifton Park, NY). B) Calculated root length densities by soil depth for simulated root profiles. C) Best Linear Unbiased Estimators for total root length (m) between DX and SX hybrids by root size class (SC) when 8 leaves were fully expanded (supplementary Table 1). D-F) Best Linear Unbiased Estimators for root systems width measured using X-rays PSC technology for root SC3 (d, 725u – 2,465u), SC4 (e, 362u – 1,232u), and SC5 (f, 181u – 616u) at stage of development V8. Predictions for root system width (W) by each depth (d) are centered. Γ functions are: W_SX,SC=3_=(0.57±0.19) × d^1.37±0.085^ × e^(0.0119±0.0007 × d)^, W_DX,SC=3_=(1.31±0.33) × d^1.143±0.06^ × e ^(0.0092±0.0005 × d)^, W_SX,SC=4_=(6.77±0.95) × d^0.68±0.03^ × e^(0.0031±0.0002 × d)^, W_DX,SC=4_=(8.80±1.08) × d^0.63±0.03^ × e^(0.0027±0.0002 × d)^, W_SX,SC=5_=(10.59±2.42) × d^0.47±0.05^ × e^(0.0019±0.00024 × d)^, W_DX,SC=5_=(8.36±1.69) × d^0.56±0.04^ × e^(0.0022±0.0002 × d)^

## Root systems architecture changed with selection

A maize hybrid set spanning a century of breeding (Table 1) was used as a case study to test the hypothesis that water capture underpins crop improvement in maize. This set comprises hybrids commercialized since 1920 that were widely adopted by farmers of the time. The sequence starts with the open pollinated Reid Yellow Dent and ends with AQUAmax^®^ drought tolerant germplasm^12^. Maize SX and DX were exposed to contrasting water treatments and plant populations to determined genetic gain in water uptake and yield. Root architecture was measured using X-ray technology (Fig. 2). Consistent with theoretical predictions^1^ (Fig. 1a,b), we show that older DX had significantly larger root systems than modern SX for all root diameter classes (*P*< 0.05; Fig. 1c). The largest effect of long-term selection manifested on the upper soil layers (Fig. 1d-f). Root morphology was modeled using the Gamma-Ricker function. Significant differences (*P*<0.05) for roots of diameters between 725 and 2,465u (supplementary Table 1) were detected between hybrid types in function parameters indicating genotypic differences in root system width with depth.

**Table 1.** List of single and double cross hybrids and year of commercial release by experiments

| Hybrid name | Cross type | Year of commercialization | Experiment |  |  |
| --- | --- | --- | --- | --- | --- |
|  |  |  | 1 | 2 | 3 |
| 351 | Double | 1934 | x |  |  |
| 322 | Double | 1936 | x |  | x |
| 317 | Double | 1937 |  |  | x |
| 340 | Double | 1941 | x |  |  |
| 344 | Double | 1945 |  |  | x |
| 352 | Double | 1946 | x | x |  |
| 347 | Double | 1950 |  |  | x |
| 301B | Double | 1952 | x | x |  |
| 3394 | Single | 1991 | x |  | x |
| 33G26 | Single | 1998 |  |  | x |
| 33P67 | Single | 1999 | x |  |  |
| 34G13 | Single | 2000 |  |  | x |
| 33R77 | Single | 2001 |  |  | x |
| 33D11 | Single | 2005 | x | x |  |
| 35A52 | Single | 2010 | x |  |  |
| P1151HR | Single | 2011 | x | x |  |

**Figure 2.**
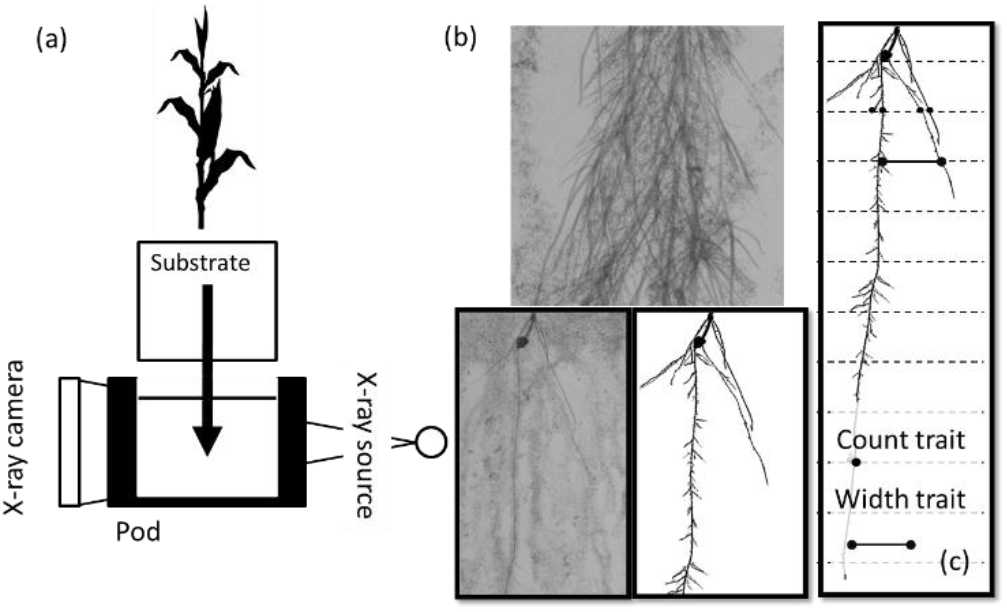
Low intensity X-ray phenotyping a) schematic of system, b) example for single image and composite, and c) illustration of count traits by depth, and width of root system.

Following principles of Darwinian agriculture^5^, observed differences between DX and SX in total root length (Fig. 1c) and width (Fig. 1d-f) could have been caused by plant-to-plant variation in root size due to genetic segregation and intraspecific competition^13^. Significant plant-to-plant variation was detected for root traits, which were measurable in total root length (*P*<0.05), root system width (*P*<0.05), and density (*P*<0.05). However, root system width is an indicator of the outer bound of root occupancy of a given volume of soil but not how effectively this volume is explored by the root system. The root length to width ratio (LWR) provides a metric to assess plausible changes from DX to SX in their capacity to explore occupied volumes. The LWR calculated from total root length and width, and for root classes 3, 4 and 5 (supplementary Table 1), are 0.17, 0.08, 0.04 cm cm^−2^ for DX, and 0.17, 0.08, 0.03 cm cm^−2^ for SX. We show that neither the allometry between root size classes nor the efficiency by which roots explore an occupied volume have changed between the DX and SX breeding eras.

## Water uptake remained unchanged over eras of breeding

Results from root morphology and water uptake experiments indicates that selection improved root system efficiency but not water capture. SX had a smaller RSA per plant when measured in growth chambers, but they captured the same volume of water as the DX (Fig. 1c,d; Fig. 3c,d) despite the similar leaf area (Table 2). While patterns of water use differed between hybrid type (*P*<0.05; Fig. 3c,d), the capacity to capture water from the soil, estimated by the change in water content between 18 and 74 days after planting, was similar for SX and DX and plant populations (Fig. 3c,d). SX and DX used water at rates of 2.7 and 2.6 mm d^−1^, and 2.8 and 2.9 mm d^−1^ when grown under low and high population, respectively (Fig. 3c,d). Differences among hybrid types occurred during the grain filling period possibly due to the capacity of modern hybrids to maintain the leaf area under stress^9^. However, data from a companion experiment showed no differences (*P*(χ^2^)>0.2)) between hybrid groups when irrigation was applied pre or post flowering. Dividing rate of water use by the average Amax (Table 2), an estimator of plant canopy size, and by TRL, an estimator of root system size^15^, here we show that DX have lower root systems efficiency than SX (0.00012 d m^−1^ vs. 0.00016 d m^−1^).

**Table 2.** Best Linear Unbiased Estimators and standard errors for main effect for crop and plant traits measured in Chile under two plant populations (3 and 10 pl m^−2^) and irrigation treatments (WD1=621 and WD2=408 mm) for double (DX) and single (SX) cross hybrids. Amax and TLN are the size of the largest leaf and the total number of leaves on the main stem, ASI is anthesis-silking interval, Scatter is the percent of the ear cob covered with non-contiguous kernels.

| Population | Location | Hybrid Group | Scatter | TLN | Amax | Time to Anthesis | ASI |
| --- | --- | --- | --- | --- | --- | --- | --- |
| (pl m<sup>-2</sup>) |  |  | (%) | (count) | (cm<sup>2</sup>) | (d) | (d) |
| 3 | WD1 | DX | 8.0±2.3 | 20.0±0.6 | 974±26 | 73.89±0.86 | 0.37±0.78 |
|  |  | SX | 3.5±1.5 | 19.4±0.5 | 1062±27 | 70.92±1.17 | -1.25±0.94 |
|  | WD2 | DX | 27.1±2.3 | 20.2±0.5 | 924±25 | 72.38±0.54 | 4.82±1.04 |
|  |  | SX | 4.5 ±1.5 | 19.4±0.5 | 1023±26 | 71.44±0.96 | -0.38±1.17 |
| 10 | WD1 | DX | 11.0±2.3 | 19.6±0.5 | 964±25 | 72.00±0.60 | 1.87±0.78 |
|  |  | SX | 5.2 ±1.5 | 19.3±0.5 | 1023±26 | 69.75±1.00 | -0.81±0.94 |
|  | WD2 | DX | 44.0±2.8 | 20.0±0.5 | 888±25 | 72.44±0.62 | 4.69±1.67 |
|  |  | SX | 7.1 ±2.2 | 19.5±0.5 | 957±26 | 72.06±1.01 | 0.56±1.75 |

**Figure 3.**
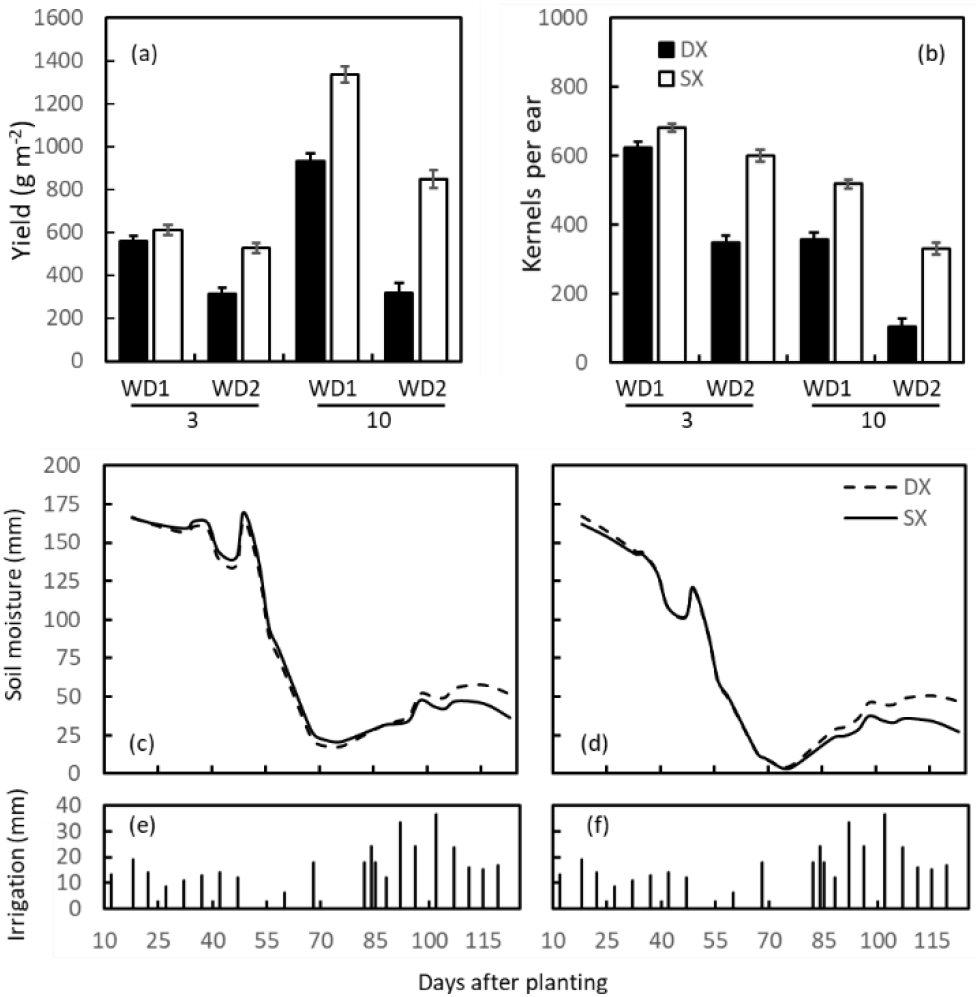
Best Linear Unbiased Estimators for contrasts between double cross (DX) and single cross (SX) maize hybrids grown in Chile for kernels per ear (a), yield (b, g m^−2^), grown under 3 pl m^−2^, and 10 pl m^−2^ and two irrigation regimes (WD1=621 and WD2=408 mm), and temporal dynamics of plant available soil water (mm) measured in WD2 in 1 m soil column and 3 pl m^−2^ (c), and 10 pl m^−2^ (d). Irrigation amounts displayed in panels e and f.

The difference in soil moisture between low and high plant populations were 17±2.8 for DX and 13±2.8 mm for SX when the soil moisture reached a minimum value. This result suggests that plant population is the main control of root occupancy and water capture, and that there was enough water in the soil column to quantify differences in soil water capture due to variation in RSA if differences were present (Fig. 3c,d). At low plant populations, between 17 and 20 mm of water was measurable in the soil. This water could have been utilized by the hybrid group with larger root systems or canopies. However, soil water content was not significantly different between the DX and SX groups when the soil moisture was at the minimum under high density (−0.8±3.8 mm, Fig. 3c,d). No differences were observed despite DX presumably having larger root systems based on the X-ray study (Fig. 1c-f).

## Yield improvement due to reproductive resilience

In contrast with results shown for water capture, yield and yield components were significantly higher for SX than DX across treatments (*P*<0.001). Yield of DX decreased with increasing barrenness, and barrenness increased with increasing stress (Table 2; Fig. 3a,b). Yield of both DX and SX decreased with decreasing kernels per ear (Fig. 3a,b). Kernels per ear decreased with increasing anthesis-silking interval (ASI) (*y*=528(±48)-67.26(±19.03)•*x*, *r*=0.81). The lower absolute correlation between kernels per ear and ASI relative to yield and ASI (|*r*|=0.81 vs. |*r*|=0.58) is associated with compensation between kernel weight and number within fertile ears^7,16^. The high association between barrenness and ASI for DX indicates that protandry induced by water deficit^7,17^ was a major driver underpinning a reduction in kernels per ear. Because both SX and DX groups reached anthesis at the same time for an irrigation and planting density treatment (Table 2), and with the same soil water content (Fig. 3c,d), we can rule out that differences in stress were due to timing of reproductive stages and soil water content. The higher ASI and barrenness observed for DX than for SX indicates that protandry for DX was long enough to miss at least part of the pollination window^7^. In addition, water deficit caused significant reductions in kernels per ear, which were larger in DX than SX (Fig. 3a), which implies differences between hybrid types in tolerance to water deficit beyond those explained by protandry alone (Fig. 3a,b; Table 2). Significant differences between populations and hybrid types in yield and yield components indicates variation in stress tolerance unrelated to water capture (Table 2; Fig.3 a,b), such as synchrony of pollination^17^, reduced sensitivity of silk elongation to drought^7,17^, and maintenance of carbon metabolism^18^.

## Yield gain from iterative genetic and agronomy optimization

We conclude that selection did not operate to increase water capture per plant and that the higher reproductive resilience in SX is not a consequence of improved water capture as postulated before^1,2^, but that yield improvement in maize is related to improved water capture due to higher planting rates that translate into higher aerial mass and yield. The differential response measured as ASI, kernels per ear and yield between SX and DX when exposed to the same level of water stress (Fig. 3c,d) provide unequivocal evidence that genetic improvement of yield precedes changes in RSA. We propose a non-dichotomous view, whereby selection for yield led to improvements in reproductive resilience, which in turn enabled changes in the structure of the plant community (Fig. 4). Changes in agronomic practices such as plant population could have led to changes in optimal root architecture, which further contributed to exposing genetic variation for RSA traits and genetic gain for yield. RSA adapted to increasingly crowded stands by decreasing the root system angle, increasing the efficiency of water uptake and reproductive resilience through shifts in C allocation. The lower total root length, higher occupancy of small roots, equal or higher total plant leaf area, and constant water uptake, suggests that SX have higher root system efficiencies measured on a per root length basis than DX. The observation that both SX and DX capture the same amount of soil water at low plant population suggests that genetic improvement operated towards optimizing RSA for improved efficiency of water capture. The reduction in RSA width, rather than being a cause of improved water capture as proposed^1,2^, is a contributor to improved root system efficiency and stress tolerance through shifts in C allocation to the ear and increased water capture through increased plant population. The hypothesis that RSA could explain interactive effects of genotype by environment (GxE) on yield^1^ was conceptually flawed because selection operated to reduce GxE in temperate maize^19^, and improved reproductive resilience is not conducive to the emergence of GxE patterns^20^.

**Figure 4.**
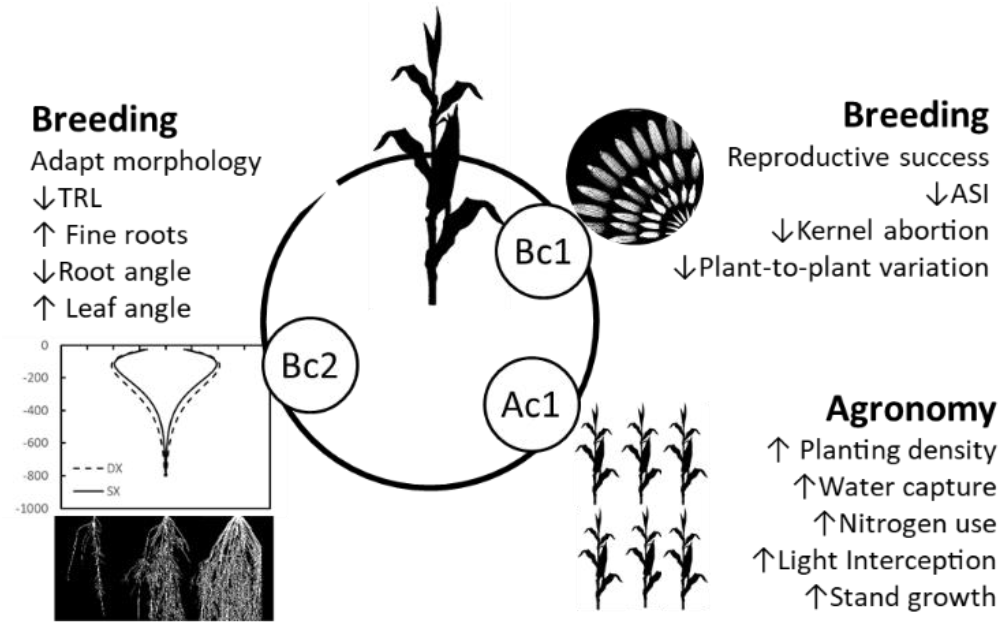
New framework for effects of selection for yield on iterative changes in plant reproductive success through resilience and root architecture co-evolving with community structure as modified by agronomic practices. ASI: anthesis-silking interval, TRL: total root length. Bc and Ac denote breeding and agronomy cycles of evaluation and optimization, with numeral indicating the position in the sequence.

The feedback between genetics and agronomy, and the evidence that the impact of root phenomics and selection on yield within breeding programs have been slow^21^, questions the feasibility of ideotype breeding in maize for root systems as proposed before^22,23^. Improved phenotyping capabilities as shown here and elsewhere^24,25,26^ can help accelerate the impact of root biology on yield improvement, but there are limitations on the speed at which one can integrate root phenotypes within breeding programs due the sequential and iterative nature of co-selection and adaptation^11^. Exploring adjacent spaces in the adaptation landscape^2^, whereby shifts in traits are tested as hypotheses within the genotype-by-management systems, can be a more productive approach to accelerate yield improvement. With a clear definition of breeding objectives and precision phenomics, prediction methodologies that integrate quantitative genetics and agronomy models^12,20,27^ offer a path to maximize the use of biological knowledge^24,25,28,29^ to accelerate genetic gain for yield can C sequestration^30^

## Materials and Methods

### Characterizing RSA in controlled environments

Root phenomics were conducted on the ERA hybrids (Table 1) at Phenotype Screening Corporation (PSC) in Knoxville, TN (experiment 1). Plants were grown in hydroponic conditions using a modified Hoagland solution (241 ppm N, 10.5 ppm P, 170 ppm K, 30 ppm Ca, 55 ppm Mg, 64.5 ppm S, 0.032 ppm B, 0.12 ppm Cu, 13 ppm Fe, 0.88 ppm Mn, 0.025 ppm Mo, 0.767 ppm Zn). Maize seeds were pregerminated and transplanted after 6 days. Phenotyping was conducted at stages V6 and V8 for root attributes and plant height to the highest fully formed collar at the vegetative development stage. Each plant container was made of fused expanded polystyrene with internal dimensions of 1000 × 45 × 200 mm. The container walls are gas permeable and allow gas exchange throughout the depth of the container. The containers were filled with expanded polystyrene beads (Alliance Foam Technologies, Centralia, MO, T180F, 1.5 pounds/cubic inch) as the growth substrate. Each container was placed in structural pods that can hold eight plants. The dripper assembly system for each container consisted of four equally spaced pressure-compensated dripper heads (Netafim Irrigation Inc., Fresno, CA, 01WPCJL2-B, 0.5GPH) operating on a 20/270 seconds on/off cycle at approximately 0.4 gallons/hour. A bank of metal halide lamps provided 400 umol m^−2^ s^−2^ illumination on a 14/10 day/night cycle. Temperature regime was 35°C/ 24°C for the day/night cycle.

A custom X-ray system developed by PSC was used to image roots growing in polystyrene containers (Fig 2). The expanded polystyrene containers are nearly transparent in the images at the X-ray energy used (25KV, 800uA.) Once placed in the X-ray chamber, a computer-controlled positioner moved the plant vertically and horizontally in predetermined steps to capture eighty 5cm × 5cm high-resolution X-ray images covering the entire one-meter deep root system. An X-ray imaging system is conceptually similar to a pin-hole camera-based system. The X-ray beam began as a point source and spread out as a cone beam. The exposure time of each X-ray images was approximately 400 milliseconds. The optical resolution of the system was approximately 58 microns. The resulting images were gray-scale images with denser and thicker root-tissue being a dark gray to black and very fine diameter root-tissue being a light gray.

Each root system was imaged two times – once at the V6 developmental stage and once at the V8 stage. Images were analyzed using RhizoTraits, version 1, a custom software developed by PSC to extract root traits from X-ray images. RhizoTraits is built off ImageJ^31^. Eighty high-resolution X-ray images were combined to create a composite image for analysis of the whole root system (7,526 pixels by 18,194 pixels, 137MP). A PSC proprietary stochastic-based segmentation algorithm was used to identify root tissue within the images. Quantitative root traits are extracted from the images and for this experiment included i.) total root length, and ii.) root system width, at each of 40 transects separated by 25 mm (supplementary Table 2, Fig. 2). Analyses were conducted for 5 root diameter size classes (supplementary Table 2; Fig. 2).

### Soil water uptake contrasting hybrid and plant populations

Previous experiments that have evaluated effects of breeding on drought tolerance measured soil moisture content at high plant populations, which may have induced plant-to-plant variation thus overestimating the role of population uniformity in water uptake. A field experiment was conducted in Viluco, Chile using the ERA hybrids^9,12^ (Table 1) to test the effects of breeding era, plant population, and water stress on soil water uptake (experiment 2). The experiment included four hybrids from the SX and DX breeding eras (Table 1), which were replicated 8 times in a split-plot design with density as the main plot, irrigation as the sub-plot, and hybrid as the sub-sub plot. Two irrigation treatments were imposed: low (high) water deficit treatments received 408 (621) mm of water for both high (10 pl/m^2^) and low (3 pl/m^2^) planting density treatment using drip tapes installed 20 cm belowground. The experiment was planted on November 7^th^, 2014 and it was harvested on April 1^st^, 2015. Soil moisture content was monitored using Time Domain Reflectometry (TDR) technology (TRIME-PICO IPH/T3, IMKO Micromodultechnik GmbH, Germany). Access tubes were installed to 1m where a rock river bed was reached. In addition to soil moisture measurements, kernel number, ear length and kernel area per ear was measured using imaging^11^, plant height, leaf number and size of the ear leaf was measured for 2 plants per plot. Time to flowering was measured for 10 individual plants daily. For analyses purposes, plants that did not flower after 86 days were assigned a value of 87 days. Leaf area was estimated by length, width and a 0.79 multiplier. The ratio between water use to leaf area per plant was used to calculate root systems efficiency^17^. Total number of leaves and leaf area of the largest plant were measured as estimators of plant leaf area^32^. Flowering notes and proportion of barren plants were recorded for 10 plants. Yield was measured using imaging methods calibrated for the location^11^.

### Field water uptake for SX and DX hybrids

A field experiment was conducted in Woodland, CA under two irrigation treatments, 8 hybrids (Table 1) of similar flowering maturity but spanning 50 years of commercial breeding, in two replications (experiment 3). The experiment was located on a Yolo soil with soil water holding capacity between 0.13 and 0.16 cm^3^cm^−3^ and no known impedance to root growth^4^. The hybrids were planted on March 27^th^, 2003 in 6 m long by 1.5 m wide plots. Plant population was 8.8 pl/m^2^. Soil moisture probes (DIVINER2000, Sentek Sensor Technologies, Australia) were installed in each experimental plot. Irrigation management was such to enable monitoring patterns of soil water uptake pre and post flowering. Irrigation was applied through drip tape installed 10 cm deep in raised planting beds. Soil water monitoring was conducted in two replications for plots subject to preflowering stress and one replication in plots subjected to postflowering stress. The average flowering date was 6/25/2003 for the group of DX hybrids, and 6/26/2003 for the SX hybrids.

## Statistical analyses

Total root length and plant height from experiment 1 were modeled within a linear mixed effect model framework with the objective to test cross type contrasts between DX and SX hybrids, with named hybrids (genotype, GE) considered as samples taken from a broader population of hybrid class,

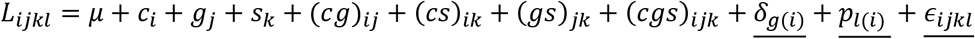

where *L_ijkl_* is total root length for plant *l* within cross type *i* and GE *g*(*i*), growth stage *j* and size class *k*. In this model, cross type (DX or SX, *c_i_*), growth stage (*g_j_*), size class (*s_k_*) and all two-way and three-way interactions were considered as fixed effects; GE (*δ*_*g*(*i*)_) and plant (*p*_*l*(*i*)_) served as random effects. The residual term is 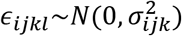 which means that for each level of cross type, growth stage and size class, a unique variance component 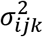 is fitted in the mixed model. For the plant height variable, a linear mixed effect model was fitted,

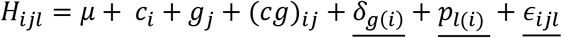

where *H_ijl_* represents plant height for cross type *i* with plant *l* at growth stage *j*. Other notations are the same as described for total root length. Cross type, growth stage and their interaction were considered fixed effects; GE and plant served as random effects. The residual term of this model is 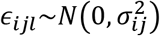, where 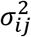 represents the residual variance term among plants for cross type *i* at growth stage *j*, allowing each level of cross type and growth stage to have a unique residual variance component.

To test for heterogeneity of plant-to-plant variation between SX and DX hybrids under different growth stages or root size classes, likelihood ratio method was applied using two nested mixed effects models. For the plant height trait, cross type fixed effect and GE (nested in cross type) random effect are included in both models.

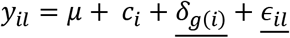

For the full model (M1), the residual variance parameter depends on the level of cross type, 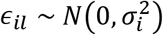. For the reduced model (M2), a single residual variance parameter is used for all observations, *ϵ_il_* ~ *N*(0, *σ*^2^). The p-value of likelihood ratio test was calculated as,

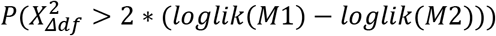

where Δ*df* = *df*(*M*1) – *df*(*M*2).

A similar approach was implemented to test plant-to-plant variation among cross types for root level traits. For a given growth stage, the full model (M1),

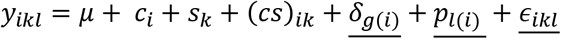

where, cross type, size class and its interaction were considered fixed effects. GE and plant were considered as random effects. For root level traits models, both the plant id random effect and residual contribute to plant-to-plant variation. In the full model (M1), the variance parameter of plant id random effect *p*_*l*(*i*)_ depends on the level of cross type, 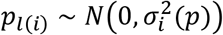. And the residual variance parameter depends on the specific level of cross type and root size class, 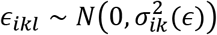.

In contrast, the reduced model (M2) considered the plant id variance parameter the same for both cross types *p*_*l*(*i*)_ ~ *N*(0, *σ*^2^(*p*)) and the residual parameter dependent only on the root class level instead of both cross type and root class, 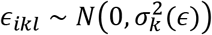.

The models to test plant-to-plant variation for traits that vary with depth (d) conditional to growth stage, M1 and M2 were extended to include the variable depth and interaction with cross type, size class as fixed effects:

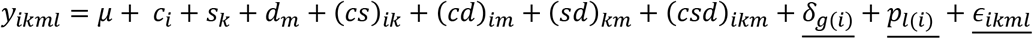

where GE and plant were considered as random effects. Just like the models of root-level traits, M1 has plant-level variance components depended on cross type, 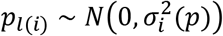, and residual variance parameter depended on cross type and root size 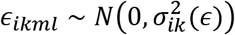. For the reduced model, both variance parameters did not vary between cross types, *p*_*l*(*i*)_ ~ *N*(0, *σ*^2^(*p*)) and 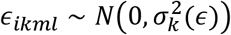.

If the underlying factor is root size class, to test the heterogeneity of plant-to-plant variation between SX and DX for two growth stages, similar method can be applied. In this case, size class was replaced with growth stage in the previous models.

The width of the root system was modeled as a function of depth using a non-linear mixed effects model, with underlying nonlinear function as Gamma-Ricker function:

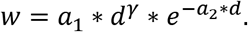

where width (*w*) is set as the response variable and depth (*d*) as the explanatory variable. For all three parameters *a*_1_, *a*_2_ and *γ*, growth stage (*j*), cross type (DX or SX, *i*) and their interaction were considered fixed effects, and GE and plant (*l*) as random effects. Taking *γ* as an example, the mixed effect model is

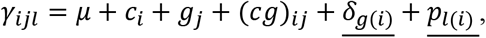

where the GE random effect 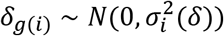 and the plant random effect 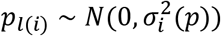. Parameters and fitted curves were estimated for each root size class (3, 4, 5).

Reproductive and vegetative traits from experiment 2 were analyzed within a generalized linear mixed effect model framework to test for differences between cross type DX and SX. For the traits with continuous numeric values, a Gaussian model with identity link was used. For the traits with count or fraction values (e.g. proportion of barren plants), a binomial model with logistic link was used. In the generalized linear mixed effects model, cross type, plant population, location and their interactions were considered fixed effects, while field spatial factor defined as row and columns, and named hybrid nested in cross type (Table 1) were considered random effects. Proportion of barren plants was modeled as,

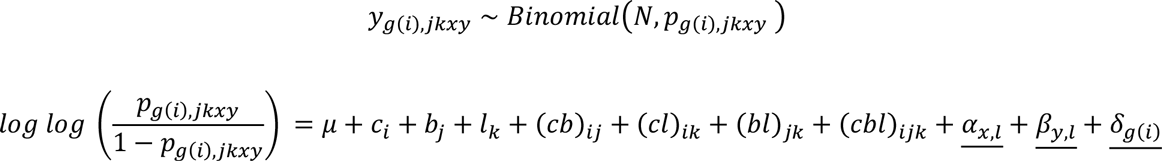

where *c_i_* is cross type (DX or SX) effect, *b_j_* is plant population effect, *l_k_* is irrigation treatment effect, (*cb*)*_ij_*, (*cl*)*_ik_*, (*bl*)*_jk_*, (*cbl*)*_ijk_* are the two factor and three factor interaction effects between cross type, population, and irrigation, *α_x,l_* and *β_y,l_* are row and column effects at each irrigation treatment with 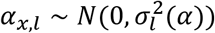 and 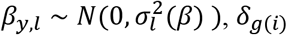 is the named hybrid random effect with 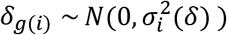 and 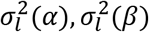 and 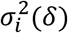 are variance parameters for the three random effects in the model.

A Generalized additive model with integrated smoothness was applied to analyze the effect of cross type, plant population and total depth (800 or 1000 mm) on the temporal dynamics of soil water content for experiment 2. The dependent variable *y* was total available soil water (mm) and the independent variable was days after planting (*x*),

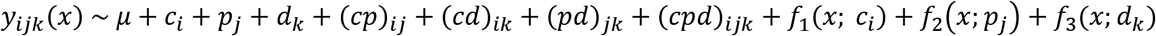

where hybrid cross (DX or SX) (*c_i_*), plant population (*p_j_*), total depth (*d_k_*) and their interactions served as the parametric terms. The functions *f*_1_(*x*; *c_i_*), *f*_2_(*x*; *p_j_*), and *f*_3_(*x*; *d_k_*) are the smoothing terms by cross type, plant population and total depth, respectively. Cubic regression spline basis with dimensions of 20 were used to fit the smoothing function *f*_1_, *f*_2_ and *f*_3_. All the nonparametric smoothing terms estimated here were centered at 0. In this model, each parametric term represents the overall magnitude of certain fixed effect, while, each smoothing term represents the pattern of the curve under each specific level of the corresponding factor.

Under each irrigation treatment, a linear mixed effect model was used to model the dynamics of soil water content at Woodland, CA measured in experiment 3. In this model, the response is soil water content; fixed effects included cross type (DX or SX), time (days after first measurement) and their interactions; random effects included named hybrid, replicates and interaction between named hybrid and time, and interaction between replicates and time. Here, factor variable named hybrid was nested within cross type. From the fitted model, we obtained the estimates of the intercept and slope for each level of cross type. Chi-square tests were conducted to evaluate the significance of cross type effect on the dynamic of water content.

All the linear mixed models and generalized linear mixed models were estimated using Asreml version 3^33^. The nonlinear mixed effect models were fitted using R package “nlme” version 3.1-144^34^. The generalized additive models were fitted using the R package “mgcv”^35^.

## Acknowledgements

This work was partially supported by the United States Department of Agriculture National Institute of Food and Agriculture grant number 2019-67019-29401.

## Author contributions

C.M. and M.C conceived the research and wrote the paper. D.McD executed the phenotyping in controlled environments and conducted the image analyses. H.P., R.C. and G.G contributed to write the paper. A.S., C.G., M.C., and C.M. conducted the field experiments. Y.F., T.T., M.C. and C.M conducted the statistical analyses.

## References

1. Hammer, G. L. et al. Can changes in canopy and/or root system architecture explain historical maize yield trends in the US Corn Belt? Crop Sci. 49, 299–312 (2009).

2. Messina, C. D., Podlich, D., Dong, Z., Samples, M. & Cooper, M. Yield–trait performance landscapes: from theory to application in breeding maize for drought tolerance. J. Exp. Bot. 62, 855–868 (2011).

3. York, L. M, Galindo-Castañeda, T., Schussler, J. R. & Lynch J. P. Evolution of US maize (*Zea mays* L.) root architectural and anatomical phenes over the past 100 years corresponds to increased tolerance of nitrogen stress. J. Exp. Bot. 66, 2347–2358 (2016).

4. Reyes, A. et al. Soil water capture trends over 50 years of single-cross maize (*Zea mays* L.) breeding in the US corn-belt. J. Exp. Bot. 66, 7339–7346 (2015).

5. Denison, R. F., Kiers, E. T. & West, S. A. Darwinian agriculture: when can humans find solutions beyond the reach of natural selection? Q. Rev. Biol 78, 145–168 (2003).

6. Edmeades, G. O., Hernandez, J. B. M. & Bello, S. Causes for silk delay in a lowland tropical maize population. Crop Sci. 33, 1029–1035 (1993).

7. Messina, C. D. et al. On the dynamic determinants of reproductive failure under drought in maize. in silico Plants 1(1) diz003 (2019).

8. Duvick, D. N. & Cassman, K. G. Post–Green revolution trends in yield potential of temperate maize in the north-central United States. Crop Sci. 39, 1622–1630 (1999).

9. Duvick, D. N. The contribution of breeding to yield advances in maize (*Zea mays* L.). Adv. Agron. 86, 83–145 (2005).

10. Assefa, Y. et al. Analysis of long-term study indicates both agronomic optimal plant density and increase maize yield per plant contributed to yield gain. Sci. Rep. 8, 4937 (2018).

11. Cooper, M. et al. Predicting the future of plant breeding: complementing empirical evaluation with genetic prediction. Crop Pasture Sci. 65, 311–336 (2014).

12. Cooper, M. et al. Integrating Genetic Gain and Gap Analysis to predict improvements in crop productivity. Crop Sci. 60, 582–604 (2020).

13. Daynard, T. B. & Muldoon, J. F. Plant-to-plant variability of maize plants grown at different densities. Can. J. Plant Sci. 63, 45–59 (1983).

14. Barker, T. et al. Improving drought tolerance in maize. Plant Breed. Rev. 25, 173–253 (2005).

15. van Oosterom, E. J. et al. Hybrid variation for root system efficiency in maize: potential links to drought adaptation. Funct. Plant Biol. 43, 502–511 (2016).

16. Borrás, L., Slafer, G. A. & Otegui, M. E. Seed dry weight response to source–sink manipulations in wheat, maize and soybean: a quantitative reappraisal. Field Crops Res. 86, 131–146 (2004).

17. Fuad Hassan, A., Tardieu, F. & Turc, O. Drought induced changes in anthesis-silking interval are related to silk expansion: a spatio-temporal growth analysis in maize plants subjected to soil water deficit. Plant Cell Environ. 31, 1349–1360 (2008).

18. Zinselmeier, C., Jeong, B. & Boyer, J. S. Starch and the control of kernel number in maize at low water potentials. Plant Physiol. 121, 25–35 (1999).

19. Gage, J. L. et al. The effect of artificial selection on phenotypic plasticity in maize. Nat. Commun. 8, 1348 (2017).

20. Messina, C. D. et al. Leveraging biological insight and environmental variation to improve phenotypic prediction: Integrating crop growth models (CGM) with whole genome prediction (WGP). Eur. J. Agron. 100, 151–162 (2018).

21. Tracy, S. R., Nagel, K. A., Postma, J. A., Fassbender, H. & Wasson, A. Crop improvement from phenotyping roots: Highlights reveal expanding opportunities. Trends Plant Sci. 25, 105–118 (2020).

22. Lynch, J. P. Steep, cheap and deep: an ideotype to optimize water and N acquisition by maize root systems. Ann. Bot., 112, 347–357 (2013).

23. Meister, R., Rajani, M. S., Ruzicka, D. & Schachtman, D. P. Challenges of modifying root traits in crops for agriculture. Trends Plant Sci. 19, 779–788 (2014).

24. Zhu, J., Ingram, P. A, Benfey, P. N. & Elich, T. From lab to field, new approaches to phenotyping root systems architecture. Curr. Opin. Plant Biol. 14, 310–317 (2011).

25. Kuijken, R. C. P., van Eeuwijk, F. A., Marcelis, L. F. M. & Bouwmeester, H. J. Root phenotyping: from component trait in the lab to breeding. J. Exp. Bot. 66, 5389–5401 (2015).

26. Atkinson, J. A., Pound, M. P., Bennett, M. J. & Wells, D. M. Uncovering the hidden half of plants using new advances in root phenotyping. Curr. Opin. Biotech. 55, 1–8 (2019).

27. Peng, B. K. et al. Advancing multiscale crop modeling for agricultural climate change adaptation assessment. Nature Plants 6, 338–348 (2020).

28. Messina, C., Cooper, M., Reynolds, M. & Hammer, G. Crop science: A foundation for advancing predictive agriculture. Crop Sci. 60, 544–546 (2020).

29. Kremling, K. et al. Dysregulation of expression correlates with rare-allele burden and fitness loss in maize. Nature 555, 520–523 (2018).

30. Kell, D. B. Large-scale sequestration of atmospheric carbon via plant roots in natural and agricultural ecosystems: why and how. Phil. Trans. R. Soc. B 367, 1589–1597 (2012).

31. Schneider, C. A., Rasband, W. S. & Eliceiri, K. W. NIH Image to ImageJ: 25 years of image analysis. Nature Methods 9, 671–675 (2012).

32. Soufizadeh, S. et al. Modelling the nitrogen dynamics of maize crops – Enhancing the APSIM maize model. Eur. J. of Agron. 100, 118–131 (2018).

33. Gilmour, A. R., Gogel, B. J., Cullis, B. R. & Thompson, R. ASReml user guide release 3.0 VSN International Ltd, Hemel Hempstead, HP1 1ES, UK (2009).

34. Pinheiro, J., Bates, D., DebRoy, S. & Sarkar, D. R. Core Team. nlme: Linear and nonlinear mixed effects models. R package version 3.1-144 (2020).

35. Wood, S. N. Fast stable restricted maximum likelihood and marginal likelihood estimation of semiparametric generalized linear models. J. R. Stat. Soc. B 73, 3–36 (2011).

