## supplementary table 1 for "Reproductive resilience but not root architecture underpin yield improvement in maize (*Zea mays* L.)"

**Supplementary data**

Supplementary Table 1. Plant and root traits measured in controlled environments. Root traits derived from features extracted from images.

| | |
|---|---|
| Measured Plant Height (mm) | height from the top of container to the leaf collar line of the last fully expanded leaf at the vegetative stage of measurement. |
| Size Class (SC) | SC1 2,900um - 9,860um<br>SC2 1,450um - 4,930um<br>SC3 725um - 2,465um<br>SC4 362um - 1,232um<br>SC5 181um - 616um |
| TRL (m) | total root length in meters of all root segments within the defined size class. |
| WidthAtDepth (mm) | width of the root system at the defined transect depth for roots in the defined size class. |
| CountDensity (#/mm2) | number of roots crossing the plane of the defined transect. The area of the plane is given by the cross-sectional area of the container used and the measured width of the root system at the defined depth. |
